# The efficacy of a 14-day modified quadruple therapy containing amoxicillin, tetracycline and high dose metronidazole and proton-pump inhibitors as an empirical third line *helicobacter pylori* eradication treatment in Taiwan

**DOI:** 10.1101/327981

**Authors:** Hsiang Tso Huang, Chih-Ming Liang, Chen-Hsiang Lee, Wei-Chen Tai, Cheng-Kun Wu, Shih-Cheng Yang, Chih-Fang Huang, Chien-Hua Chiu, Kai-Lung Tsai, Meng-wei Chang, Hsin-Ming Wang, Keng-Liang Wu, Ping-I Hsu, Deng-Chyang Wu, Seng-Kee Chuah, on behalf of Taiwan Acid-Related Disease (TARD) Study Group

## Abstract

The antibiotics resistances to amoxicilln, tetracycline was low in Taiwan even after multiple *H. pylori* treatment failures and high dose metronidazole could overcome antibiotics resistance. In real world practice, susceptibility-guided treatments are not widely available. Therefore, we assessed the efficacy of 14-day modified quadruple therapy containing amoxicillin, tetracycline and high dose metronidazole and PPI as an empirical third-line rescue *H. pylori* treatment. This study was conducted by analyzing 70 consecutive prospectively registered patients who failed two times *H. pylori* eradication. All of them received endoscopy for *H. pylori* culture. Seven patients were lost to follow up. They were then treated according to the antibiotic susceptibility testing reports (Cultured group, n=39). Those who failed *H. pylori* culture were prescribed with a modified 14-day quadruple therapy containing esomeprazole 40 mg twice daily, amoxicillin 1 g twice daily, tetracycline 500 mg four times daily and metronidazole 500 mg three times daily (empirical group, n=24). Follow-up urea breath test was performed 8 weeks later. The eradication rates attained by Cultured group and empirical group were 89.7% (95% confidence interval [CI] = 72.72%-97.11%) and 58.3% (95% CI=36.61%-77.86%), in the per protocol analysis (p=0.004); 81.4%(95% CI=66.60%-91.61%) and 51.8% (95% CI=31.9%-71.29%), in the intention-to-treat analysis (p=0.014). Culture-guided therapy was the clinical factors influencing the efficacy of *H. pylori* eradication (OR: 0.16; 95% CI: 0.04-0.60, p=0.006). In conclusion, empirical 14-day modified 3 quadruple therapy is not acceptable as an alternative third-line rescue *H. pylori* treatment arobably but the success rate of the third-line susceptibility-guided treatment was only moderate (<90%).

## Introduction

*Helicobacter pylori (H. pylori)* infection is one of the most common bacterial infections in the world and was classified as grade I carcinogen (1–3). It could slowly induce chronic gastritis, which progresses through the premalignant stages of atrophic gastritis, intestinal metaplasia, and dysplasia, and then finally to gastric cancer (4, 5). Meanwhile, successful eradication of *H pylori* has greatly reduced the recurrence of peptic ulcers (6). The worldwide standard triple therapy consisted with one proton pump inhibitor (PPI) and amoxicillin and clarithromycin, and in place of amoxicillin for who are allergic to it, metronidazole has been one alternative. However, *H pylori* may develop resistance to the prescribed antibacterial like clarithromycin as standard first line therapy and may acquire resistance by acquisition and recombination of genes from other bacteria (7). The resistance battle against *H. pylori* continue, many people still abuse self-medication of antibiotic like levofloxacin, which helps *H. pylori* to develope more drugs resistances (8–10).

After failure of second-line treatment, culture with susceptibility testing or molecular determination of genotype resistance is recommended by the Maastricht V/Florence-Consensus Report (11). The secondary resistances to clarithromycin and levofloxacin are high in patients who failed regimens containing these antibiotics (12–13) Therefore, the reuse of clarithromycin and levofloxacin empirically should be avoided in the third line regiments. On the other hand, there is a major limitation to this therapeutic line because of low sensitivity of culture-based advice. Moreover, majority of hospitals and clinics have no facility for culture with antibiotics susceptibility testing or molecular test of genotype resistance. The more effective third line regiments for *H. pylori* should be warranted in the future. The issue on the urge for an ideal empirical third line *H. pylori* rescue therapy has been raised. The Toronto consensus recommended the choice of antibiotics empirically according to medication history (14). However, earlier studies have generally shown poor eradication rates with Rifabutin based (15) or Rifaximin based (16) as third line rescue therapy of only 63-66% regardless of combination with amoxicillin, levofloxacin or clarithromycin.

In Taiwan, *H. pylori* culture with susceptibility testing or molecular determination of genotype resistance is recommended in third line rescue therapy in adherence to Taiwan Consensus Report (17). However, we may encounter patients who failed *H. pylori* culture or refuse to do further endoscopy and molecular determination of genotype resistance is not available. Liou et al reported that secondary antibiotic resistances to amoxicillin and tetracycline are low in Taiwanese patients who failed at least two eradications (18). The addition of a PPI helped overcome metronidazole resistance and double dose twice a day is recommended (19). In our previous studies, we observed that high dose metronidozole may overcome metronidazole (20). Therefore, the present study aims to evaluate the efficacy of this simple modified empirical 14-day quadruple therapy with esomeprazole, amoxicillin, metronidazole and tetracycline as the third line regiments for *H. pylori* eradication.

## Results

### Patients’ demographic and baseline characteristics

Figure 1 summarizes the patient disposition. Seven patients were lost to follow up. Therefore, 63 patients were included in the per protocol analysis. Their baseline characteristics are shown in table 1. The mean age is 58.9 ± 10.1 years. 36.5% are male patients and 63.5% are female patients. There was no significant difference between the culture group and empirical group in age, gender, smoking and alcohol habits, peptic ulcer history, and the endoscopic finding

**Figure 1.**
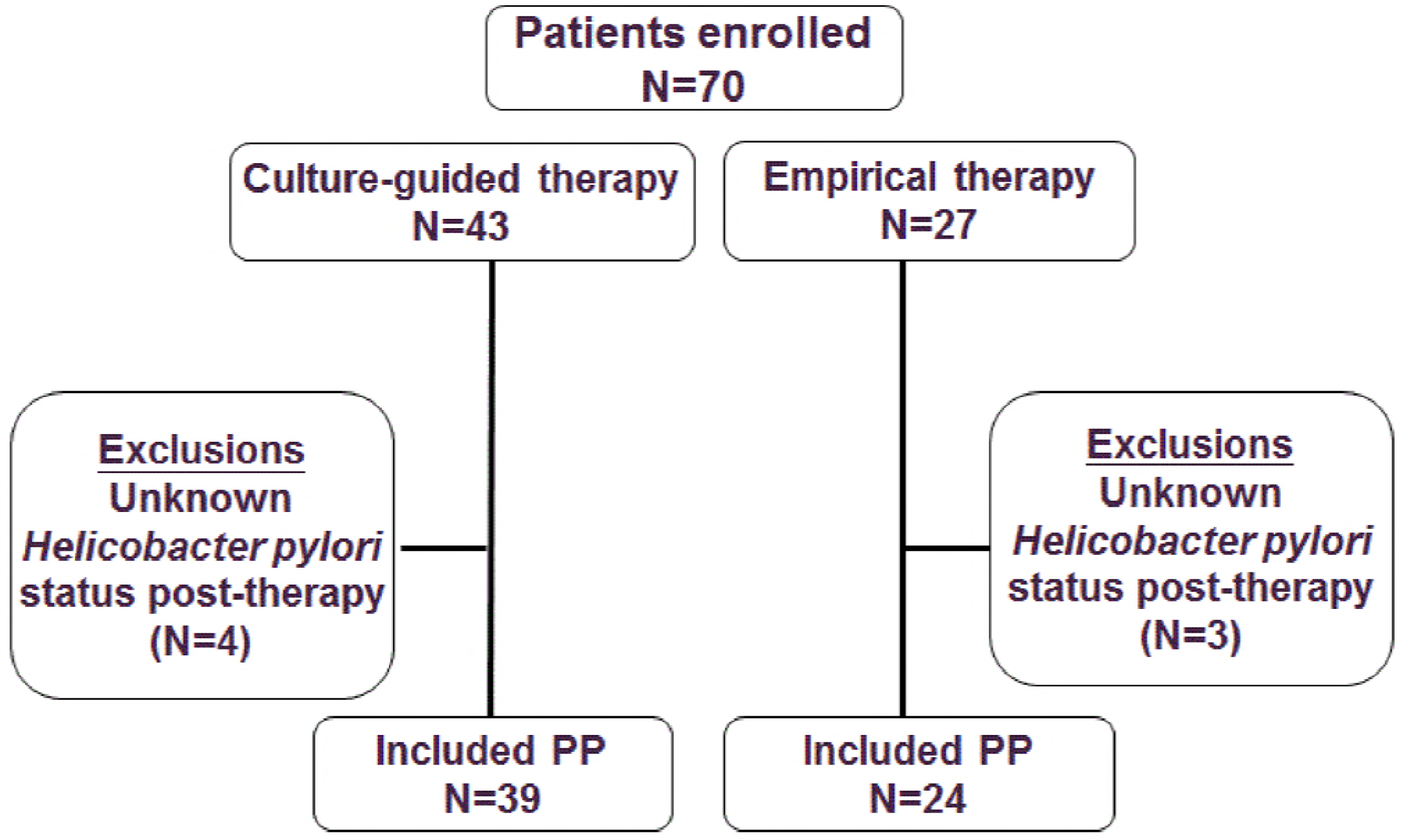
Disposition of patients.

**Table 1.** Demographic data and endoscopic appearances of the two patient groups.

| Characteristics | Culture-guided therapy<br>(n =39, %) | Empirical therapy<br>(n =24,%) | P-value |
| --- | --- | --- | --- |
| Age (year) (mean ± SD) | 58.7±10.4 | 58.3±10.0 | 0.869 |
| Gender (male/female) | 13/26(33.3/66.7) | 10/14(41.7/58.3) | 0.512 |
| Smoking | 4(10.3) | 1(4.2) | 0.075 |
| Alcohol consumption | 6(15.4) | 3(12.5) | 0.755 |
| Previous history of peptic ulcer | 18(46.2) | 13(54.2) | 0.293 |
| Endoscopic Findings |  |  |  |
| Gastritis | 15(38.5) | 12(50.0) | 0.063 |
| Gastric ulcer | 20(51.3) | 5(20.8) |  |
| Duodenal ulcer | 1(2.6) | 6(25.0) |  |
| Gastric and duodenal ulcer | 3(7.7) | 1(4.2) |  |

### Eradication rate and susceptibility of *H. Pylori* culture

Figure 2 demonstrated the antibiotics resistances after two *H. pylori* treatment failures. Clarithromycin resistance strains were found in 79.5% (31/39) of the patients. Levofloxacin-resistant strains were 94.9% (37/39). Metronidazole resistance strains were 66.7% (26/39). Amoxicillin resistance was 2.6% (1/39). No strains developed resistance to tetracycline. Table 2 demonstrates the major outcomes of eradication rate. The eradication rates attained by Cultured group and empirical group were 89.7% (95% confidence interval [CI] = 72.72%-97.11%) and 58.3% (95% CI=36.61%-77.86%), respectively in the per protocol analysis (p=0.004) 81.4% (95% CI=66.6%-91.61%) and 51.8% (95% CI=31.9%-71.29%), respectively, in the intention-to-treat analysis (p=0.014). Despite the high metronidazole resistance after two treatment failures, the eradication rate in the patients with was as high as 84% in this patient cohort (Table 3).

**Figure 2.**
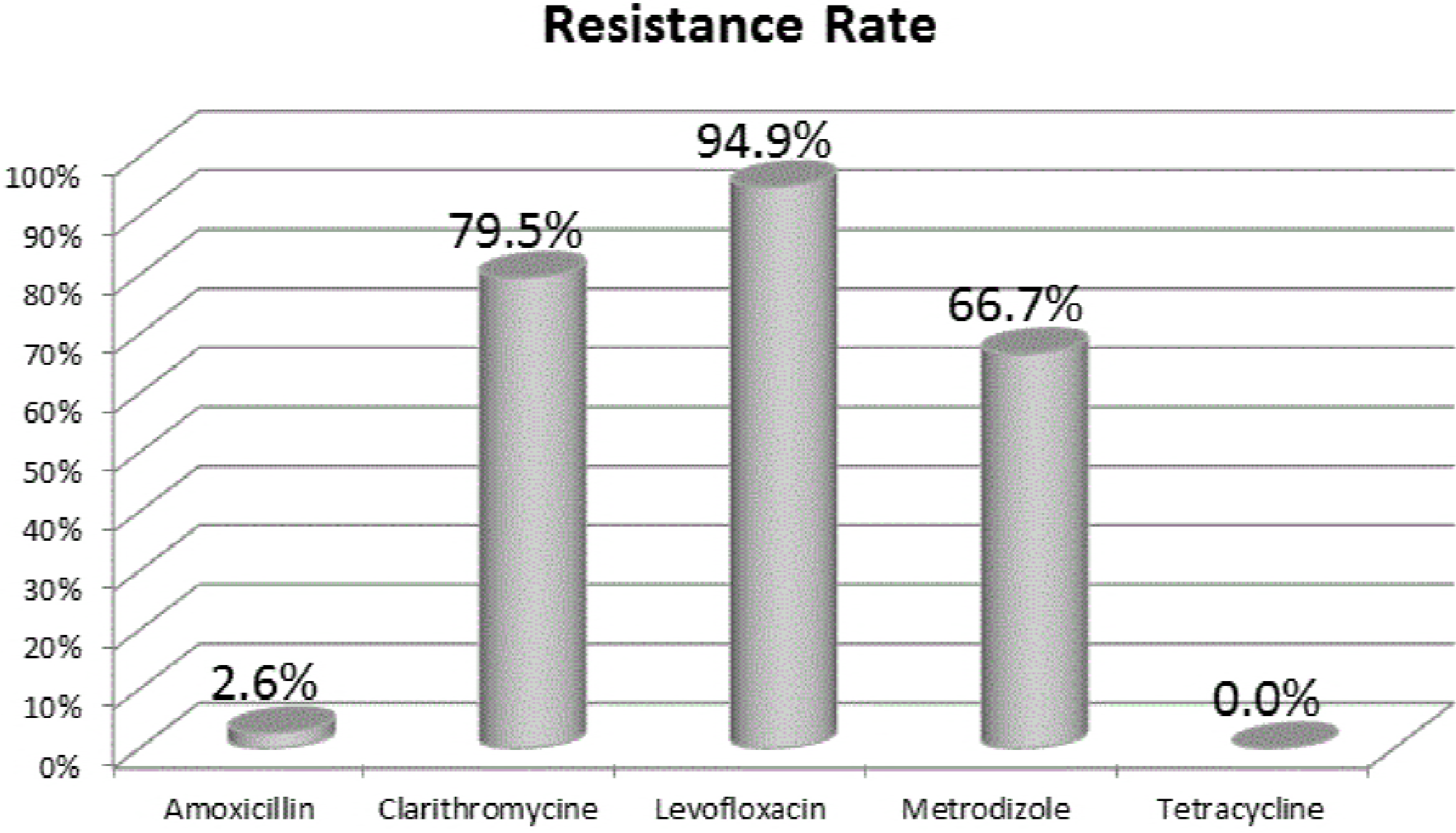
The antibiotics resistances after two *H. pylori* treatment failures.

**Table 2.** Major outcomes of eradication therapy.

|  | **Eradication rate** |  |  |
| --- | --- | --- | --- |
|  | Culture-guidedgroup | Empirical group | P-value |
| **Intention To Treat** | 81.4% (35/43) | 51.8% (14/27) | 0.014 |
| **Per-protocol** | 89.7% (35/39) | 58.3% (14/24) | 0.004 |
| **Adverse event** | 25.6% (10/39) | 29.2% (7/24) | 0.758 |
| **Compliance** | 100 % (39 /39) | 100% (24 /24) | - |

**Table 3.** Antibiotic resistance and *H. pylori* eradication rate.

| Effect of amoxicillin, tetracycline and metronidazole resistance on *Helicobacter pylori* eradication rates in the PP population |  |  |
| --- | --- | --- |
| Culture | Total | Eradication rate, N( % ) |
| AMO-S<br>TET-S<br>MET-S | 13 | 13 (100) |
| AMO-R<br>TET -S<br>MET-S | 0 | - |
| AMO-S<br>TET -R<br>MET-S | 0 | - |
| AMO-S<br>TET -S<br>MET-R | 25 | 21(84) |
| AMO-R<br>TET -R<br>MET-S | 0 | - |
| AMO-R<br>TET -S<br>MET-R | 1 | 1 (100) |
| AMO-S<br>TET -R<br>MET-R | 0 | - |
| AMO-R<br>TET -R<br>MET-R | 0 | - |
| Total | 39 | 35(89.7) |
| AMO: amoxicillin; TET: tetracycline; MET, metrodinazole; S: sensitive, R, resistant. PP: per protocol |  |  |

### Adverse events and compliance

The adverse event occurred in 17 patients (22.9%). There was no significant difference between two groups in adverse events (25.6% vs. 29.2%, p=0.758) (Table 2). The greatest side effect from the use of medications was the digestive system such as the presence of nausea, abdominal pain and constipation (Table 4). All patients completed their course of treatment, therefore, with 100% of compliance.

**Table 4.**
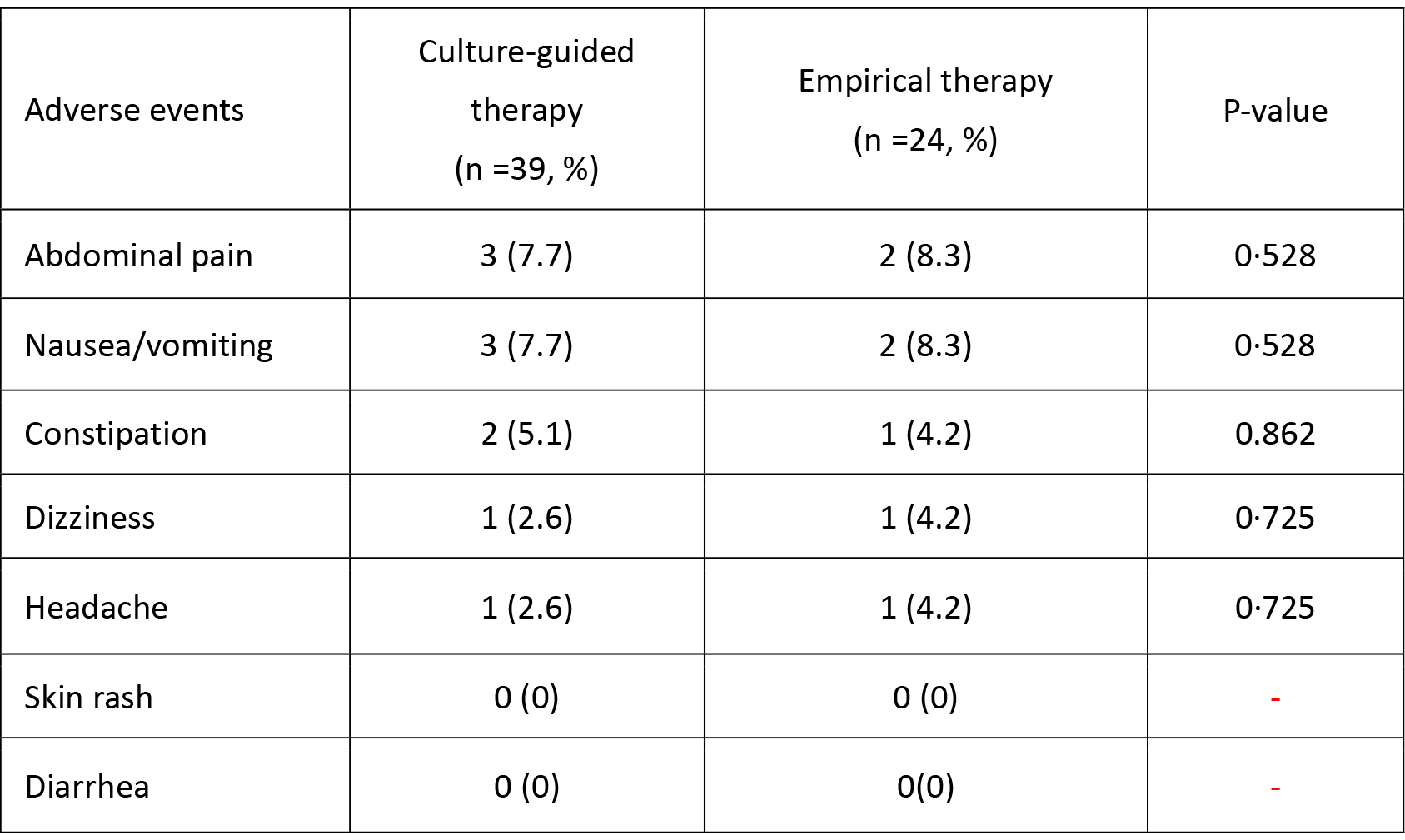
Adverse events of patients.

### Factors influenced the efficacy of anti-*H. pylori* therapy

Table 5 shows the univariate analysis of all demographic data and factors. In univariate analysis, there was only one clinical factor, culture vs. empirical therapy, with p-value 0.004. Multivariate analysis, culture vs. empirical therapy was the only clinical factor influencing *H. pylori* eradication (OR: 0.16, 95% CI: 0.04-0.60, p=0.006) (Table 6).

**Table 5.** Univariate analysis of the clinical factors influencing the efficacy of *H. pylori* eradication.

| Clinical factors |  | Eradicated |  | Not eradicated |  |  | P-value |
| --- | --- | --- | --- | --- | --- | --- | --- |
|  |  | No | % | No | % | Total |  |
| Sex | Male | 18 | 78.3 | 5 | 21.7 | 23 | 0.944 |
|  | Female | 31 | 77.5 | 9 | 22.5 | 40 |  |
| Age | <60 years | 25 | 75.8 | 8 | 24.2 | 33 | 0.686 |
|  | ≥60 years | 24 | 80.0 | 6 | 20.0 | 30 |  |
| Smoking | No | 44 | 75.9 | 14 | 24.1 | 58 | 0.460 |
|  | Yes | 5 | 100.0 | 0 | 0 | 5 |  |
| Alcohol consumption | No | 40 | 74.1 | 14 | 25.9 | 54 | 0.083 |
|  | Yes | 9 | 100.0 | 0 | 0 | 9 |  |
| Previous history of peptic ulcer | No | 25 | 75.8 | 8 | 24.2 | 33 | 0.387 |
|  | Yes | 24 | 80.0 | 6 | 20.0 | 30 |  |
| Hypertension | No | 35 | 76.1 | 11 | 23.9 | 46 | 0.595 |
|  | Yes | 14 | 82.4 | 3 | 17.6 | 17 |  |
| Diabetes Mellitus | No | 40 | 76.9 | 12 | 23.1 | 52 | 0.723 |
|  | Yes | 9 | 81.8 | 2 | 18.2 | 11 |  |
| Chronic renal failure | No | 49 | 79.0 | 13 | 21.0 | 62 | 0.059 |
|  | Yes | 0 | 0 | 1 | 100.0 | 1 |  |
| Cardiac vascular disease | No | 46 | 78.0 | 13 | 22.0 | 59 | 0.890 |
|  | Yes | 3 | 75.0 | 1 | 25.0 | 4 |  |
| Chronic hepatitis | No | 44 | 78.6 | 12 | 21.4 | 56 | 0.668 |
|  | Yes | 5 | 71.4 | 2 | 28.6 | 7 |  |
| Psychological problem | No | 47 | 77.0 | 14 | 23.0 | 61 | 0.442 |
|  | Yes | 2 | 100.0 | 0 | 0 | 2 |  |
| Hyperthyroidism | No | 48 | 77.4 | 14 | 22.6 | 62 | 0.590 |
|  | Yes | 1 | 100.0 | 0 | 0 | 1 |  |
| Culture vs, Empirical | Empirical | 14 | 58.3 | 10 | 41.7 | 24 | 0.004 |
|  | Culture-guided | 35 | 89.7 | 4 | 10.3 | 39 |  |
| Compliance | Complete treatment | 49 | 77.8 | 14 | 22.2 | 63 | - |
|  | Not complete | 0 | 0 | 0 | 0 | 0 |  |

**Table 6.** Multivariate analysis of the clinical factors influencing the efficacy of *H. pylori* eradication.

| Clinical factor | Odds ratio (95% CI) | *P*-value |
| --- | --- | --- |
| Culture guided |  |  |
| Vs. Empirical therapy | 0.16 (0.04~0.60) | 0.006 |

## Discussion

According to Maastricht V/Florence Consensus Report (11), after failure of second line therapy for eradication of *H. Pylori*, clinically, the best choice of therapy should be based on the cultivation of *helicobacter pylori* and get its susceptibility to provide a better solution for the population. However, determination of the MIC for *H. pylori* is not widely available, because it is time consuming, inconvenient and relatively expensive. Besides, the successful culture rate ranges from 75% to 90% (21, 22). Genotypic resistance-guided test is a quick and effective assay to decide the choice of antibiotics sensitive to *H. Pylori*. Liou et al (18) reported a study in the North Taiwan about genotypic resistance-guided test in third-line treatment, improving the overall eradication rate in patients who received clarithromycin-, levofloxacin- and tetracycline-based sequential therapies were 78.9% (15/19), 92.2% (47/51) and 71.4% (25/35) in strains susceptible to clarithromycin, levofloxacin and tetracycline, respectively. The eradication rate will get the optimal result in the patients who are taking the levofloxacin-based regiment if the genotype was sensitive to levofloxacin. However, in current study, the patient failing *H. pylori* eradication after using the 1st line standard triple clarithromycin-based therapy and 2nd line levofloxacin-based therapy developed the high resistant rate in clarithromycin 79.5% (31/39) and levofloxacin 94.9% (37/39). Conversely, the amoxicillin and tetracycline resistance was as low as 2.6% (1/39), and 0%. Therefore, we used a simple and modified 14-day high-dose quadruple therapy with esomeprazole, amoxicillin, metronidazole and tetracycline as the empirical third line regiment for *H. pylori* eradication for patients who failed *H. pylori* culture and refused further endoscopy procedure. There were several reasons for the empirical use of this modified quadruple empirical therapy. First, the prevalence of tetracycline and amoxicillin resistance, even in patients who have failed multiple treatments, remains low worldwide (17, 23). Our results confirmed that the prevalence of tetracycline resistance was 0% in patients failing two eradication treatments and developed 0~10% in five years’ sequential change in our previous culture report (20). Second, the impact of tetracycline and metronidazole resistance on the eradication rate remains controversial (21, 24–27). It is also because of high dose metronidazole was effective for eradication therapy, despite a relatively high resistance rate. Third, several randomized controlled trials (28, 29) compared two different durations of eradication therapy, and demonstrated that the longer duration is more effective. Therefore, we set the third line treatment course as 14 days.

In this study, per-protocol analysis, culture guided therapy achieved significantly higher eradication rate than empirical therapy (89.7% vs. 58.3%, p=0.004). This study suggests that this empirical quadruple therapy is not acceptable in third line *H. pylori* eradication therapy. However, there was no definite factor found in detail analysis to predict the successful rate, except the culture-guided vs. empirical group (p-value is 0.004). Chen et al reported a > 90 % eradication rate by using a bismuth-containing quadruple therapy with metronidazole and amoxicillin is an alternative to classical bismuth quadruple therapy for *H. pylori* third line rescue treatment (30). At this point, we wonder why reinvent the wheel as a similar regime with different doses and doing intervals ha been reported to be successful. The possible explanation could be probably related to a drug-drug interaction between amoxicillin and tetracycline which exhibit low in vitro antibiotic resistance (31). The success in Chen’s study could be due to the synergism that occurs between metronidazole and its hydroxymetabolite and between either the analog or amoxicillin/tetracycline, both of which may contribute to the efficacy of both the amoxicillin-metronidazole and tetracycline-metronidazole combinations (32). It was proposed that bacteriostatic drugs like tetracycline might interfere with the bactericidal action of penicillin (33). The bactericidal action is to inhibit cell wall formation which is dependent on how fast the bacteria are multiplying. Tetracycline may reduce the effectiveness of penicillin because it is a bacteriostatic antibiotics and can inhibit the cellular protein synthesis that is required for cell division. Eventually, susceptibility-guided treatment is still recommended for third line *H. pylori* eradication therapy as Maastricht V/Florence Consensus Report (11).

It was not so surprised that the eradication rate in the patients with metronidazole resistance was as high as 84% (21/25) as had been observed in many previous publications [20, 29, 34]. The possible explanation was environment changes in the oxygen pressure in the stomach, as metronidazole-resistant *H. pylori* isolates can become susceptible to metronidazole under low oxygen conditions in vitro (27, 35). This phenomenon of high dose metronidazole overcoming antibiotics resistance was also documented in our previous study (29, 33). In Spain, Ignasi Puig et al made a systematic review and meta-analysis on therapy based on culture-guided susceptibility (36), they found that the eradication rate of *H. pylori* was only with an average of 72%. In our study, our eradication rate was also < 90%.

In real world practice, despite the introduction of culture testing or molecular genotype resistance after failure of second-line treatment, it is not routinely available in majority of medical institutions. In this regional study, we noted that if the patients failed the clarithromycin-based standard triple therapy and levofloxacin-based therapy, the resistant strains to clarithromycin and levofloxacin were very high. Initial hypothesis was that the empirical modified 14-day high-dose quadruple therapy with amoxicillin, tetracycline, high dose metronidazole and esomeprazole, could be an alternative the third line regiment for *H. pylori* eradication. Despite that the good drug compliance and was easier to meet patients’ adherence, the eradication success rate did not fulfill our expectation probably due to the drug-drug interaction between amoxicillin and tetracycline.

There were some limitations in this study. First, the study sample size was small. Second, we were unable to turn this study into a randomized trial because we felt it was not ethical to randomize patients with multiple *H. pylori* eradication failures with available culture susceptibility results. In conclusion, empirical 14-day modified quadruple therapy is not acceptable as an alternative third-line rescue *H. pylori* treatment in spite of the low amoxicillin and tetracycline resistances probably due to the drug-drug interaction between amoxicillin and tetracycline but the success rate of the third-line susceptibility-guided treatment was only moderate (<90%). The optimal management of rescue treatments requires further research.

## Materials and methods

### Patients

A total of 70 prospective registered consecutive *H. pylori* infected patients, who failed H. pylori eradication by using the 1st line standard triple therapy and 2nd line levofloxacin-based therapy between 1, January 2015 and 31, December 2017 at Kaohsiung Chang Gung Memorial Hospital, Taiwan, were recruited in this study. Based on the international guidelines, these patients had received endoscopy for *H. pylori* culture and consequent antibiotic susceptibility testing. Seven patients were lost to follow up. All patients would be at least 18 years of age and had received endoscope examinations that showed peptic ulcers or gastritis. The confirmation of *H. pylori* eradication failure was defined as patients with either one positive 13 C-UBT or any two positive of the rapid urease test, histology and culture after second-line eradication therapy. Criteria for exclusion included (a) ingestion of antibiotics, bismuth, or PPIs within 4 weeks before; (b) patients with allergic history to the medications used; (c) patients with previous gastric surgery; (d) pregnant women. They were then treated according to the antibiotic susceptibility testing reports as 3rd line therapy (Cultured group, n=39). Those who failed *H. pylor*i culture and did not want to repeat another endoscopy for *H. pylori* culture were prescribed with a simple empirical 14-day modified quadruple therapy containing esomeprazole 40 mg twice daily, amoxicillin 1 g twice daily, tetracycline 500 mg four times daily and metronidazole 500 mg three times daily (empirical group, n=24). To assess eradication efficacy after the completion of anti-*H. pylori* therapy urea breath test was used to confirm the *H. pylori* status at 8 weeks. The absence of *H. pylori* after eradication therapy was defined as a negative result of urea breath test at 8 weeks (9, 37).

The primary outcome variables are the eradication rate, presence of adverse events, and level of patient compliance. Demographic information including age, gender, social history of smoking, alcohol consumption, previous peptic ulcer history and medical history were collected via electrical medical records. This study was approved by both the Institutional Review Board and Ethics Committee of Chang Gung Memorial Hospital, Taiwan (IRB 201800034B0).

### Culture and antimicrobial susceptibility testing

One biopsy specimen from the antrum and one from the corpus were obtained for *H. pylori* isolation using previously described culture methods (10, 11). The biopsy specimens were cultured on plates that contained Brucella chocolate agar with 7% sheep blood and were incubated for 4-5 d under micro-aerobic conditions. The minimal inhibitory concentration (MIC) was determined by the agar dilution test. The *H. pylori* strains were tested for susceptibility to amoxicillin, Clarithromycin, levofloxacin, metronidazole and tetracycline by using the E-test method (AB BIODISK, Solna, Sweden). *H. pylori* strains had MIC values of ≥ 0. 5, ≥ 1, ≥ 1, ≥ 4 and ≥ 8 mg/L, which were considered to be the resistance breakpoints for amoxicillin, Clarithromycin, levofloxacin, metronidazole and tetracycline, respectively (38)

### Statistical analysis

By using the SPSS program (Statistical Package for the Social Sciences version 23, Chicago, IL, USA), Chi-square tests with or without Yates’ correction for continuity and Fisher’s exact tests were used when appropriate to compare the major outcomes between groups. Eradication rates were analyzed by per-protocol (PP) approaches. The PP analysis excluded patients with unknown H. pylori status following therapy and those with major protocol violations. A P-value <0.05 was considered statistically significant. To determine the independent factors that affected treatment response, the clinical parameters were analyzed by univariate and multivariate analysis.

## Acknowledgement

The authors would like to acknowledge Miss Ching-Yi Lin for her assistance in this study.

## References

1. Cave DR. 1997. How is Helicobacter pylori transmitted? Gastroenterology 113 (6 Suppl): S9–14.

2. World Health Organization, International Agency for Researchon Cancer (IARC). IARC monographs on the evaluation of carcinogenic risks to humans. 1994. Schistosomes, liver flukes and Helicobacter pylori. Lyon: IARC, 61 177–240.

3. Graham DY. 1997. Helicobacter pylori infection in the pathogenesis of duodenal ulcer and gastric cancer: a model. Gastroenterology 113 1983–1991.

4. Bin Lu, Meng Li. 2014. Helicobacter pylori eradication for preventing gastric cancer. World J Gastroenterol 20 5660–5.

5. Correa P, Piazuelo MB.2012. The gastric precancerous cascade. J Dig Dis 13 2–9

6. Hopkins RJ, Girardi LS, Turney EA. 1996. Relationship between Helicobacter pylori eradication and reduced duodenal and gastric ulcer recurrence: a review. Gastroenterology 110 1244–1252

7. Tursi A, Picchio M, Elisei W. 2012. Efficacy and Tolerability of a Third-Line, Levofloxacin-Based, 10-Day Sequential Therapy in Curing Resistant Helicobacter Pylori Infection. J Gastrointestin Liver Dis 21 133–138.

8. Murakami K, Furuta T, Ando T, Nakajima T, Inui Y, Oshima T, Tomita T, Mabe K, Sasaki M, Suganuma T, Nomura H, Satoh K, Hori S, Inoue S, Tomokane T, Kudo M, Inaba T, Take S, Ohkusa T, Yamamoto S, Mizuno S, Kamoshida T, Amagai K, Iwamoto J, Miwa J, Kodama M, Okimoto T, Kato M, Asaka M; Japan GAST Study Group. 2013. Multi-center randomized controlled study to establish the standard third-line regimen for Helicobacter pylori eradication in Japan. J Gastroenterol 48 1128–1135

9. Chuah SK, Tai WC, Hsu PI, Wu DC, Wu KL, Kuo CM, Chiu YC, Hu ML, Chou YP, Kuo YH, Liang CM, Chiu KW, Hu TH. 2012. The Efficacy of Second-line Anti-Helicobacter Pylori Therapy Using an Extended 14-days levofloxacin/amoxicillin/protonpump inhibitors-A Pilot Study. Helicobacter 17 374–381

10. Chuah SK, Tai WC, Lee HC, Liang CM. 2014. Quinolone-containing therapies in the eradication of Helicobacter pylori. Biomed Res Int 2014 151543.

11. Malfertheiner P, Megraud F, O’Morain CA, Gisbert JP, Kuipers EJ, Axon AT, Bazzoli F, Gasbarrini A, Atherton J, Graham DY, Hunt R, Moayyedi P, Rokkas T, Rugge M, Selgrad M, Suerbaum S, Sugano K, El-Omar EM; European Helicobacter and Microbiota Study Group and Consensus panel. 2017. Management of Helicobacter pylori infection-the Maastricht V/Florence Consensus Report.Gut. 66 6–30.

12. Liou JM, Chang CY, Chen MJ, et al. The primary resistance of Helicobacter pylori in Taiwan after the national policy to restrict antibiotic consumption and its relation to virulence factors-Anationwidestudy. PLoS ONE. 2015;10 e0124199.

13. Chang WL, Sheu BS, Cheng HC, Yang YJ, Yang HB, Wu JJ. 2009. Resistance to metronidazole, clarithromycin and levofloxacin of Helicobacterpylori before and after clarithromycin-based therapy in Taiwan. J GastroenterolHepatol. 24 1230–1235.

14. Fallone CA, Chiba N, van Zanten SV, Fischbach L, Gisbert JP, Hunt RH, Jones NL, Render C, Leontiadis GI, Moayyedi P, Marshall JK. 2016. The Toronto Consensus for the Treatment of Helicobacter pylori Infection in Adults. Gastroenterology. 151 51–69.

15. Gisbert JP, Calvet X. 2012. Review article: rifabutin in the treatment of refractory Helicobacter pylori infection. Aliment Pharmacol Ther 35 209–221

16. Perri F, Festa V, Clemente R, Villani MR, Quitadamo M, Caruso N, Bergoli ML, Andriulli A. 2001. Randomized study of two “rescue” therapies for Helicobacter pylori-infected patients after failure of standard triple therapies. Am J Gastroenterol 96 58–62.

17. Sheu BS, Wu MS, Chiu CT, Lo JC, Wu DC, Liou JM, Wu CY, Cheng HC, Lee YC, Hsu PI, Chang CC, Chang WL, Lin JT. 2017. Consensus on the clinical management, screening-to-treat, and surveillance of Helicobacter pylori infection to improve gastric cancer control on a nationwide scale. Helicobacter. 22. doi: 10.1111/hel.12368.

18. Liou JM, Chen CC, Chang CY, Chen MJ, Fang YJ, Lee JY, Chen CC, Hsu SJ, Hsu YC, Tseng CH, Tseng PH, Chang L, Chang WH, Wang HP, Shun CT, Wu JY, Lee YC, Lin JT, Wu MS; Taiwan Helicobacter Consortium. 2013. Efficacy of genotypic resistance-guided sequential therapy in the third-line treatment of refractory Helicobacter pylori infection: a multicentre clinical trial. J Antimicrob Chemother. 68 450–456.

19. Graham DY, Lee SY. 2015. How to Effectively Use Bismuth Quadruple Therapy: The Good, the Bad, and the Ugly. Gastroenterol Clin North Am. 44 537–563.

20. Chuah SK, Liang CM, Lee CH, Chiou SS, Chiu YC, Hu ML, Wu KL, Lu LS, Chou YP, Chang KC, Kuo CH, Kuo CM, Hu TH, Tai WC. 2016. A randomized control trial comparing two levofloxacin-containing second-line therapies for Helicobacter pylori eradication. Medicine (Baltimore) 95 e3586

21. Gisbert JP.2008. "Rescue" regimens after Helicobacter pylori treatment failure. World J Gastroenterol. 14: 5385–5402.

22. Vakil N, Megraud F. 2007. Eradication therapy for Helicobacter pylori. Gastroenterology. 133 985–1001.

23. Wu IT, Chuah SK, Lee CH, Liang CM, Lu LS, Kuo YH, Yen YH, Hu ML, Chou YP, Yang SC, Kuo CM, Kuo CH, Chien CC, Chiang YS, Chiou SS, Hu TH, Tai WC. 2015. Five-year sequential changes in secondary antibiotic resistance of Helicobacter pylori in Taiwan. World J Gastroenterol. 21 10669–10674

24. Fischbach L, Evans EL. 2007. Meta-analysis: the effect of antibiotic resistance status on the efficacy of triple and quadruple first-line therapies for Helicobacter pylori. Aliment Pharmacol Ther 26 343–357.

25. Laine L, Hunt R, El-Zimaity H, Nguyen B, Osato M, Spénard J.2003. Bismuth-based quadruple therapy using a single capsule of bismuth biskalcitrate, metronidazole, and tetracycline given with omeprazole versus omeprazole, amoxicillin, and clarithromycin for eradication of Helicobacter pylori in duodenal ulcer patients: a prospective, randomized, multicenter, North American trial. Am J Gastroenterol 98 562–567

26. Nista EC, Candelli M, Cremonini F, Cazzato IA, Di Caro S, Gabrielli M, Santarelli L, Zocco MA, Ojetti V, Carloni E, Cammarota G, Gasbarrini G, Gasbarrini A. 2003. Levofloxacin-based triple therapy vs. quadruple therapy in second-line Helicobacter pylori treatment: a randomized trial. Aliment PharmacolTher 18 627–633

27. Gerrits MM, van der Wouden EJ, Bax DA, van Zwet AA, van Vliet AH, de Jong A, Kusters JG, Thijs JC, Kuipers EJ. 2004. Role of the rdxA and frxA genes in oxygen-dependent metronidazole resistance of Helicobacter pylori. J Med Microbiol 53 1123–8

28. Treiber G, Wittig J, Ammon S, Walker S, van Doorn LJ, Klotz U. 2002. Clinical outcome and influencing factors of a new short-term quadruple therapy for Helicobacter pylori eradication: a randomized controlled trial (MACLOR study). Arch Intern Med 162 153–160.

29. Kongchayanun C, Vilaichone RK, Pornthisarn B, Amornsawadwattana S, Mahachai V. 2012. Pilot studies to identify the optimum duration of concomitant Helicobacter pylori eradication therapy in Thailand. Helicobacter 17 282–5.

30. Chen Q, Zhang W, Fu Q, Liang X, Liu W, Xiao S, Lu H. 2016. Rescue Therapy for Helicobacter pylori Eradication: A Randomized Non-Inferiority Trial of Amoxicillin or Tetracycline in Bismuth Quadruple Therapy. Am J Gastroenterol 111 1736–1742.

31. Wu DC, Hsu PI, Tseng HH, Tsay FW, Lai KH, Kuo CH, Wang SW, Chen A. 2011. Helicobacter pylori infection: a randomized, controlled study comparing 2 rescue therapies after failure of standard triple therapies. Medicine (Baltimore). 90 180–185

32. Moellering RC Jr. 1983. Rationale for use of antimicrobial combinations. Am J Med 75 4–8.

33. Sorice F, Ortona L, Pizzigallo E. 1975. Further aspects of combination antibiotic therapy. Critical review and personal case studies. Minerva Med 66 2805–2822.

34. Kuo CH, Hsu PI, Kuo FC, Wang SS, Hu HM, Liu CJ, Chuah SK, Chen YH, Hsieh MC, Wu DC, Tseng HH. 2013. Comparison of 10-day bismuth quadruple therapy with high-dose metronidazole or levofloxacin for second-line Helicobacter pylori therapy: a randomized controlled trial. J Antimicrob Chemother 68 222–228.

35. Moon JY, Kim GH, You HS, Lee BE, Ryu DY, Cheong JH, Jung JI, Jeong JH, Song CS, Song GA. 2013. Levofloxacin, Metronidazole, and Lansoprazole Triple Therapy Compared to Quadruple Therapy as a Second-Line Treatment of Helicobacter pylori Infection in Korea. Gut Liver 7 406–410

36. Puig I, López-Góngora S, Calvet X, Villoria A, Baylina M, Sanchez-Delgado J, Suarez D, García-Hernando V, Gisbert JP. 2016. Systematic review: third-linesusceptibility-guided treatment for Helicobacter pylori infection. Therap Adv Gastroenterol. 9 437–448.

37. Liang CM, Cheng JW, Kuo CM, Chang KC, Wu KL, Tai WC1, Chiu KW, Chiou SS, Lin MT, Hu TH, Chuah SK.2014. Levofloxacin-containing second-line anti-Helicobacter pylori eradication in Taiwanese real-world practice. Biomed J. 37 326–330

38. Chuah SK, Hsu PI, Chang KC, Chiu YC, Wu KL, Chou YP, Hu ML, Tai WC, Chiu KW, Chiou SS, Wu DC, Hu TH. 2012. Randomized comparison of two on-bismuth-containing second-line rescue therapies for Helicobacter pylori. Helicobacter. 17 216–223.

